# EloR interacts with the lytic transglycosylase MltG at midcell in *Streptococcus pneumoniae* R6

**DOI:** 10.1101/2020.12.18.423453

**Authors:** Anja Ruud Winther, Morten Kjos, Marie Leangen Herigstad, Leiv Sigve Håvarstein, Daniel Straume

## Abstract

The ellipsoid shape of *Streptococcus pneumoniae* is determined by the synchronized actions of the elongasome and the divisome, which have the task of creating a protective layer of peptidoglycan (PG) enveloping the cell membrane. The elongasome is necessary for expanding PG in the longitudinal direction whereas the divisome synthesizes the PG that divides one cell into two. Although there is still little knowledge about how these two modes of PG synthesis are coordinated, it was recently discovered that two RNA-binding proteins called EloR and KhpA are part of a novel regulatory pathway controlling elongation in *S. pneumoniae*. EloR and KhpA form a complex that work closely with the Ser/Thr kinase StkP to regulate cell elongation. Here, we have further explored how this regulation occur. EloR/KhpA is found at midcell, a localization fully dependent on EloR. Using a bacterial two-hybrid assay we probed EloR against several elongasome proteins and found an interaction with the lytic transglycosylase homolog MltG. By using EloR as bait in immunoprecipitation assays, MltG was pulled down confirming that they are part of the same protein complex. Fluorescent microscopy demonstrated that the Jag domain of EloR is essential for EloR’s midcell localization and its interaction with MltG. Since MltG is found at midcell independent of EloR, our results suggest that MltG is responsible for recruitment of the EloR/KhpA complex to the division zone to regulate cell elongation.

**Importance:** Bacterial cell division has been a successful target for antimicrobial agents for decades. How different pathogens regulate cell division is, however, poorly understood. To fully exploit the potential for future antibiotics targeting cell division, we need to understand the details of how the bacteria regulate and construct cell wall during this process. Here we have revealed that the newly identified EloR/KhpA complex, regulating cell elongation in *S. pneumoniae*, forms a complex with the essential peptidoglycan transglycosylase MltG at midcell. EloR, KhpA and MltG are conserved among many bacterial species and the EloR/KhpA/MltG regulatory pathway is most likely a common mechanism employed by many Gram-positive bacteria to coordinate cell elongation and septation.

## Introduction

In order to multiply, a bacterial cell splits into two daughter cells in an intricate process involving chromosome replication and segregation, production of new cell membrane, and synthesis of new cell wall. *Streptococcus pneumoniae* is a Gram-positive species, meaning it produces a thick cell wall that surrounds and protects the cell. The major component of the cell wall is peptidoglycan (PG) which is made up of chains of polysaccharides that are cross linked with short peptide bridges. The polysaccharides consist of alternating molecules of N-acetylglucosamine (GlcNAc) and N-acetylmuramic acid (MurNAc). The cross links are made between pentapeptides attached to MurNAc (1).

*S. pneumoniae* has an ellipsoid shape resulting from synthesis of the PG layer by two protein complexes – the elongasome and the divisome (2, 3). As the name suggests, the elongasome is responsible for producing PG in the peripheral direction, creating the elongated shape of pneumococci. The divisome, on the other hand, is responsible for synthesizing the septal disc that divides one cell into two. The precursor for PG is made inside the pneumococcal cell, transported to the outside and incorporated into the growing PG through transglycosylation (TG) and transpeptidation (TP) reactions (1, 4). One group of enzymes performing this incorporation are the penicillin binding proteins known as PBPs. *S. pneumoniae* has six PBPs, three class A PBPs (PBP1a, PBP1b, PBP2a) that harbor both TG and TP activity, two class B PBPs (PBP2b, PBP2x) that only harbor TP activity, and PBP3, a D,D-carboxypeptidase whose activity affects the amount of cross links in PG by removing the terminal D-Ala residues of pentapeptides (5–7). It is widely acknowledged that PBP2b is an essential part of the elongasome and PBP2x is an essential part of the divisome (8–10). The Shape Elongation Division and Sporulation (SEDS) proteins RodA and FtsW have emerged as the main TG enzymes during PG production, working alongside the TP enzymes PBP2b and PBP2x, respectively. These essential protein pairs (PBP2b/RodA and PBP2x/FtsW) are the core PG polymerizing units in *S. pneumoniae* (3, 11). The discovery that SEDS proteins are the primary TG enzymes in PG synthesis, has prompted reassessment of the role class A PBPs have in PG synthesis. Rather than being essential in building the primary PG, recent data strongly indicate that the class A PBPs are essential for maturation of newly synthesized PG, e.g. filling in gaps or mistakes left by the divisome and possibly the elongasome (12, 13). Other proteins considered to be part of the elongasome and divisome are found to be important for scaffolding, localization and regulation of PG production. One newly identified member of the elongasome is the membrane bound lytic transglycosylase MltG (14). MltG consists of a cytosolic domain, a transmembrane α-helix, and an extracellular catalytic domain. several lines of evidence support that MltG is part of the elongasome: Cells depleted of MltG have reduced length, sfGFP-MltG co-localizes with elongasome proteins throughout the cell cycle, and suppressor mutations have been found in *mltG* upon deletion of the essential elongasome transpeptidase *pbp2b* demonstrating a functional link between these genes (15). The specific function of MltG in cell division is still unknown, but it has been proposed to release PG strands synthesized by PBP1a for cross-linking by RodA/PBP2b (15).

A particularly interesting aspect of cell wall synthesis is how PG production is regulated. By tracking the incorporation of new PG material using super-resolution fluorescence microscopy, both the elongasome and divisome PG synthesis machineries have been shown to be organized in regularly spaced nodes in pneumococci (16), and the cells seem to elongate a short time period before septal PG synthesis is initiated (9, 17). Although several proteins are known to be involved in regulation of PG synthesis (18–24), there is little knowledge about how the temporal and spatial control of elongation and division is achieved. The eukaryotic-type Ser/Thr kinase StkP appears to play a key role in coordinating these two events (25–27). StkP phosphorylates and thereby modulates the activity of several cell division proteins, i.e. DivIVA, GpsB, MapZ, MurC, MacP and EloR (also known as Jag/KhpB) (18, 21, 24, 28–31). DivIVA and its paralogue GpsB together with StkP are important for tuning septal and peripheral PG synthesis (18, 32), and a phosphorylated MacP regulates the function of the class A PBP PBP2a (24). Furthermore, phosphorylation of MapZ (scaffolding protein for FtsZ) has been shown to be important for FtsZ ring constriction and splitting, while the effect of MurC (UDP‐N‐acetylmuramoyl L‐alanine ligase catalyzing addition of alanine to UDP-MurNAc at an early step of the PG synthesis pathway) phosphorylation is still unclear. StkP is also important for the localization of PBP2x through interaction between StkP’s PASTA domains and the pedestal and/or the transpeptidase domain of PBP2x (33).

StkP-dependent phosphorylation of EloR has also been shown to be essential in regulation of cell elongation in *S. pneumoniae* (19, 21). EloR (short for <u>Elo</u>ngasome <u>R</u>egulating protein) is conserved in a range of Gram-positive genera, including *Streptococcus*, *Bacillus*, *Clostridium*, *Listeria*, *Enterococcus*, *Lactobacillus* and *Lactococcus*. The protein is composed of three domains: (i) an N-terminal Jag domain, (ii) a KH-II domain and (iii) an R3H domain at its C-terminal end (Fig. 1A), but no transmembrane segment. The KH-II and R3H are both single stranded nucleic acid binding domains that usually bind RNAs, while the Jag domain has an unknown function. EloR interacts with another RNA-binding protein called KhpA (composed of one KH-II domain).

**Figure 1.**
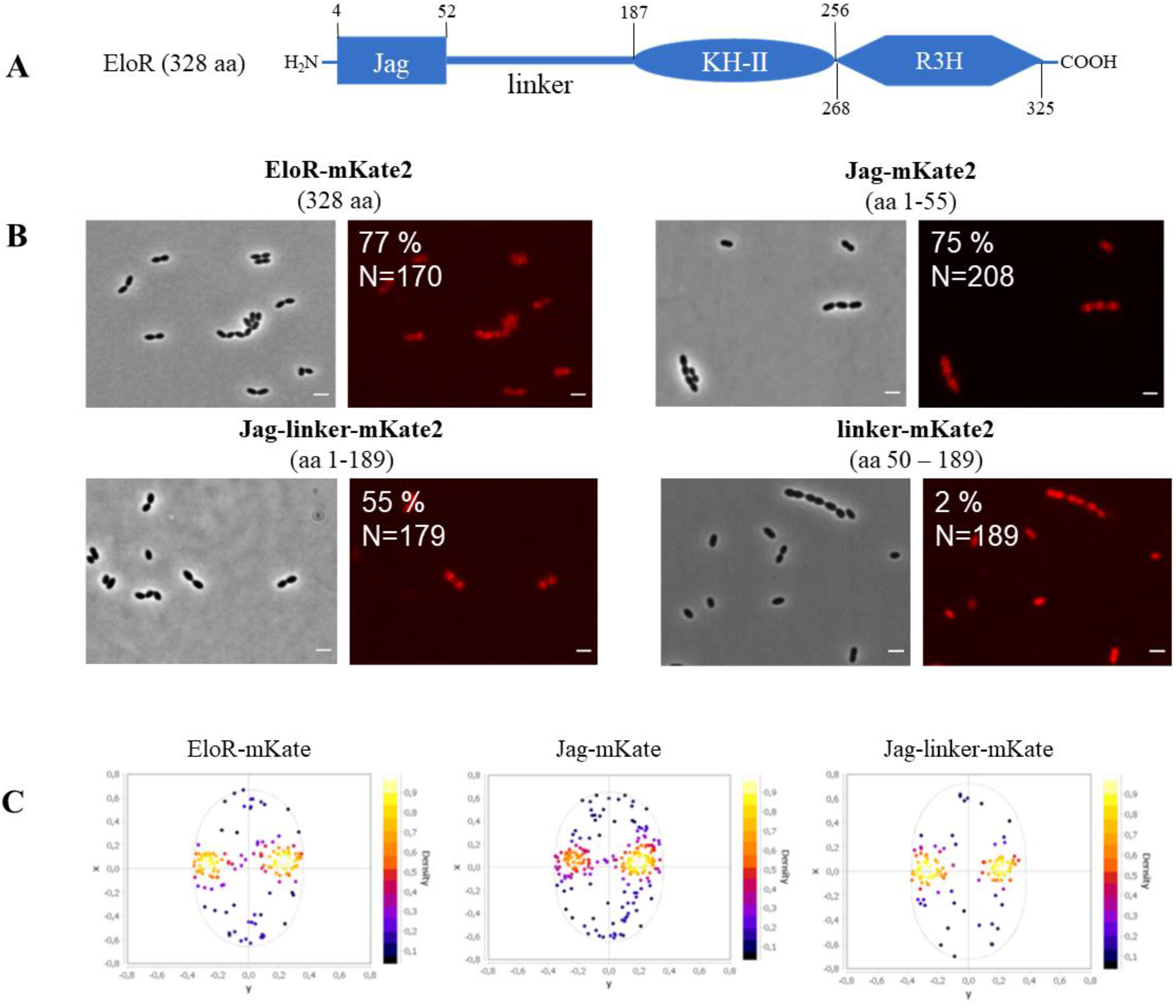
The Jag domain directs EloR to midcell. A) Schematic representation of EloR, including predicted domain borders. B) Micrographs showing the subcellular localization of EloR-mKate2 (AW407), Jag-mKate2 (AW408), Jag-linker-mKate2 (AW409), and linker-mKate2 (AW410). The numbers above the micrographs indicate the amino acids of EloR utilized in the different constructs. The numbers in the FL images indicate the percentage of cells that displayed midcell localization of mKate2. The number of cells included in the analyses are also indicated. Scale bars are 2 μm. C) Analysis of subcellular localization. Detection of fluorescent maximum signals for EloR-mKate, Jag-mKate, and Jag-linker-mKate were performed and plotted using MicrobeJ.

If the EloR/KhpA complex is broken, cells become shorter, consistent with loss of elongasome function, and are no longer dependent upon the essential PBP2b/RodA pair (19, 21). Point mutations inactivating the RNA-binding domains of EloR suggest that phosphorylation of EloR by StkP leads to release of bound RNA. This stimulates cell elongation in an unknown fashion (19). Interestingly, knockdown of *eloR* and *khpA* expression in the rod-shaped *Lactobacillus plantarum* also resulting in shortening of cells, suggesting a conserved role of these proteins in regulating cell elongation (34). EloR and KhpA localize to the division zone of *S. pneumoniae*. While midcell localization of KhpA depends on its interaction with the KH-II domain of EloR, it is not known what directs EloR to midcell.

In this study, we employed fluorescence microscopy and protein-protein interaction assays to further explore the EloR-mediated regulation of cell elongation in *S. pneumoniae*. We show that the Jag-domain of EloR is critical for the midcell localization of this protein. Furthermore, EloR was shown to interact with the elongasome protein MltG via its Jag domain, suggesting a role of MltG in positioning EloR at midcell.

## Results and discussion

### The Jag domain is solely responsible for recruiting EloR to midcell

EloR consists of an N-terminal Jag domain and two C-terminal RNA-binding domains, KH-II and R3H (Figure 1A). We and others have previously shown that EloR localizes to the division zone where it forms a complex with KhpA (20, 21). While KhpA depends on its interaction with EloR in order to localize to the division zone, it is not known how EloR finds midcell. We hypothesized that EloR must form interaction(s) with other elongasome proteins in order to localize correctly. The Jag domain is connected to the KH-II domain by a large linker region (134 amino acids long) with an unknown structure and function. Since the KH-II and R3H domains bind RNA, our rationale was that the Jag-linker part of EloR would be important for its subcellular localization. We tested this by fusing full length EloR, the Jag domain, the linker region, and the Jag-linker domains to the far-red fluorescent protein mKate2, creating the strains AW407, AW408, AW410, and AW409, respectively. These fusions were expressed ectopically from an inducible promoter using the ComRS system (35). The native *eloR* gene was kept unchanged in the genome. When inducer (ComS) was supplied to the growth medium, we saw, as expected, that full length EloR fused with mKate2 (EloR-mKate2) was concentrated at midcell for 77 % of the cells investigated (Figure 1B and C). It should be noted that we observe a background signal from the cytoplasm in most cells, suggesting that not all EloR-proteins are midcell localized. We also found that the Jag-mKate2 and the Jag-linker-mKate2 fusions concentrated at midcell for 75 % and 55 % of the cells, respectively (Figure 1B). The linker-mKate2 fusion, on the other hand, did not localize to midcell (Figure 1B), as only two percent of the linker-mKate2 cells investigated displayed a midcell fluorescent signal.

These results clearly suggest that the Jag domain is solely responsible for localizing EloR to midcell. The 3D structure of the Jag domain has been solved for EloR in *Clostridium symbiosum* (PDB number 3GKU). It has a β-α-β-β fold with the α-helix laying on top of a three-stranded β-sheet. The conserved motif KKGFLG is found in the loop connecting the β2 and β3-strands (Supplemental Figure S1A). The same is true for the predicted structure of EloR from *S. pneumoniae* (Supplemental Figure S1B). We hypothesized that the conserved region (KKGFLG) could be involved in a protein-protein interaction possibly important for EloR localization. However, substitutions of several residues (K36A, K37A, F39A, and L40M) in this motif did not abrogate the midcell localization of EloR (Supplemental Figure S2).

### The StkP-mediated phosphorylation is not critical for EloR-localization

The results above clearly suggest that the Jag domain targets EloR to the division zone independently of the linker domain. Nevertheless, the fact that the conserved threonine (threonine 89 in *S. pneumoniae*) phosphorylated by StkP to modulate EloR activity is located in the linker domain suggests that the linker could be involved in conformational rearrangements of the EloR protein between the active and inactive form. The StkP kinase is located at midcell in *S. pneumoniae* and to explore whether this protein or the phosphorylation state affected the localization of EloR (36, 37), we analyzed the localization of EloR-mKate2 in a genetic background lacking *stkP* (Δ*stkP*::janus). However, this demonstrated that StkP is not the reason why EloR-mKate2 is concentrated at midcell, since the protein retains its localization in cells lacking *stkP* (Figure 2A).

**Figure 2.**
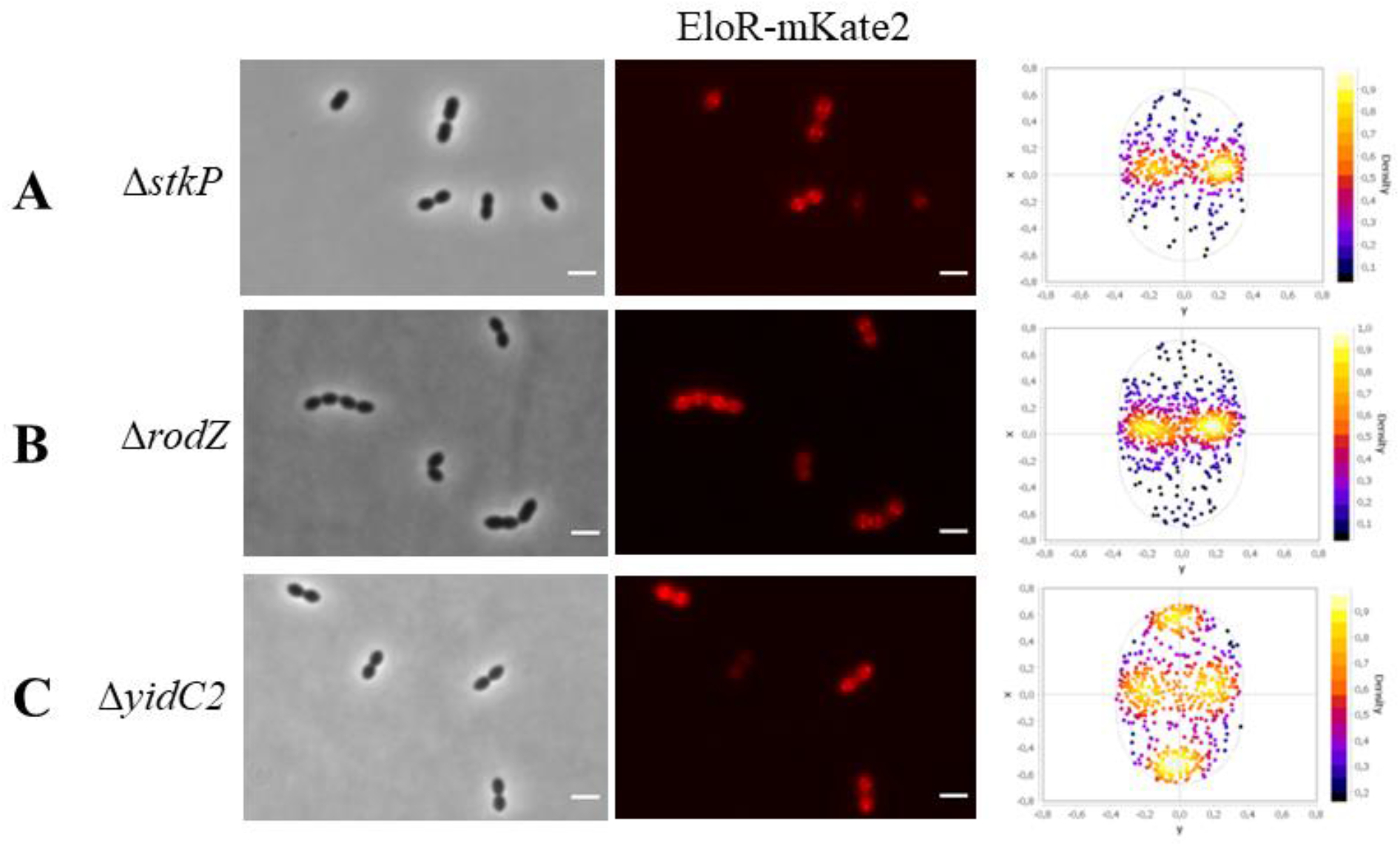
EloR-mKate2 localization in different genetic backgrounds. Localization of EloR-mKate2 in A) Δs*tkP* (N = 1155), B) Δ*rodZ* (N = 1681) and C) Δ*yidC2* (N= 1280) mutants. Scale bars are 2 μm.

Based on our results, the linker region is not crucial for recruiting EloR to midcell. Interestingly, when aligning the amino acid sequences of EloR homologues from different Gram-positive species, the length of the linker region varies from approximately 135 amino acid residues in *S. pneumoniae* to approximately 10 residues in *Bacillus subtilis* (Supplemental Figure S3). The reason for these variations is not clear, but if the linker domain is involved in protein-protein interactions the larger linker region in the pneumococcal EloR could accommodate for more interaction partners and hence more regulatory possibilities. Why pneumococci would need this is not clear. The structural fold of the EloR linker in *S. pneumoniae* is unknown, making it particularly interesting for future studies to explore its 3D structure and the rearrangements occurring between phosphorylated EloR and the non-phosphorylated form.

### EloR interacts with several proteins known to be part of the elongasome

EloR has been shown to interact with the mid-cell localized proteins StkP and KhpA, but these interactions did not affect the localization of EloR. In order to investigate how EloR localizes at midcell and to understand its regulatory function in cell elongation, we wanted to explore what other protein interactions EloR forms in addition to the one with KhpA and StkP. We screened our Bacterial Two-Hybrid (BACTH) library in *Escherichia coli* for possible interaction partners for EloR. The BACTH assay is based on blue (positive) and white (negative) color selection, where the blue color comes from cleavage of X-gal in the medium by β-galactosidase. Briefly, the two proteins that are tested for interaction are fused to either the T18 or T25 domain. If an interaction between the two proteins occurs, T18 and T25 reconstitute an adenylate cyclase producing cAMP which induces expression of β-galactosidase (38). EloR was probed against a range of known cell division proteins, namely PBP2b, RodA, RodZ, MreC, MreD, CozE, and MltG (Figure 3). We also tested YidC2 whose gene shares operon with *eloR*. YidC2 is an insertase that assists in insertion of membrane proteins into the lipid bilayer, working together with the SecYEG translocon, the signal recognition particle (SRP) and the SRP-receptor FtsY (39, 40). The presence of *yidC2* and *eloR* in one operon seems to be conserved in several species, e.g. *S. pneumoniae*, *S. mitis*, *S. oralis*, *B. subtilis*, and *Listeria monocytogenes*, indicating a functional link between the two.

**Figure 3.**
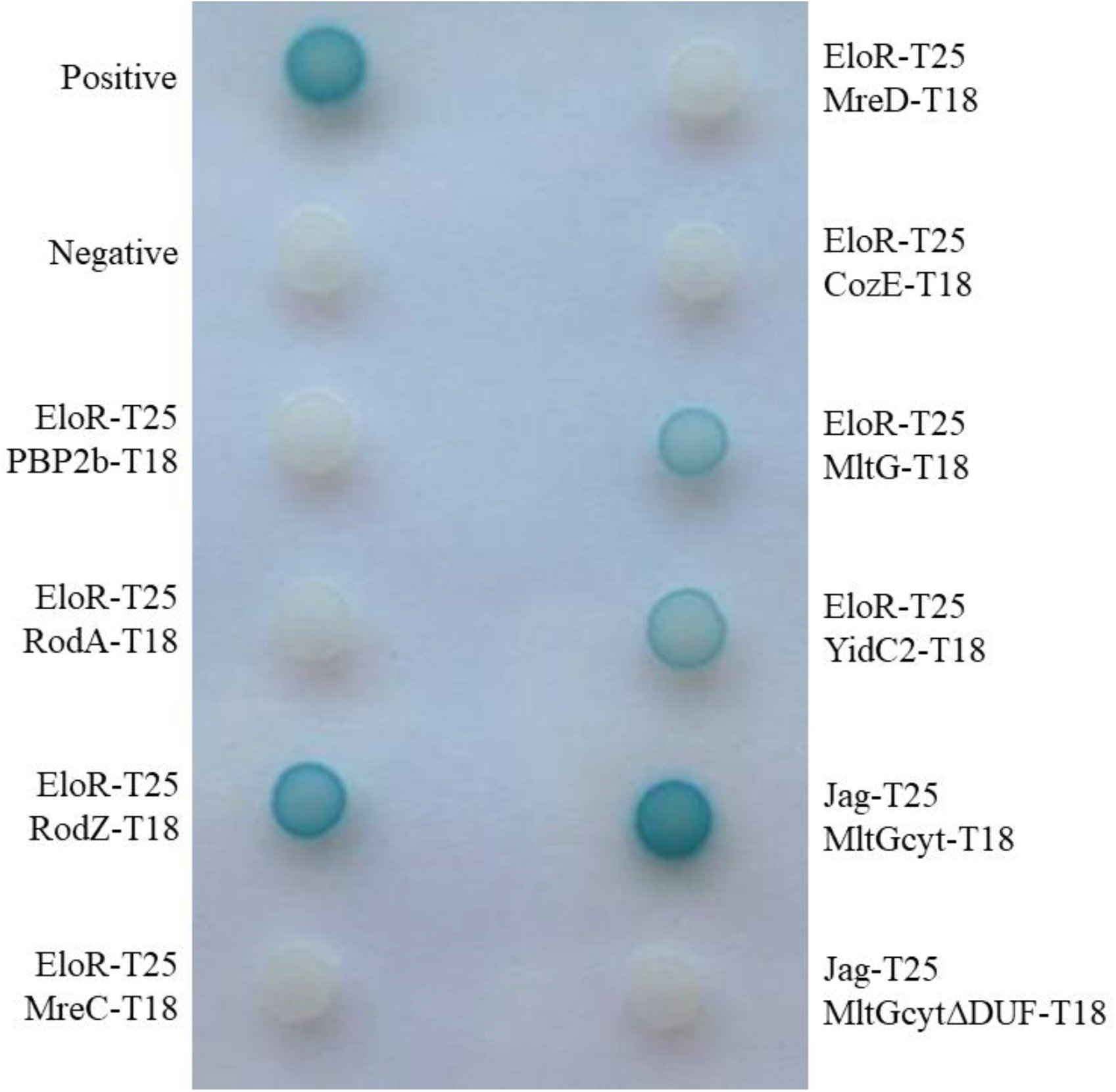
Bacterial two hybrid assay probing EloR against other elongasome proteins. PBP2b, RodA, MreC, MreD, and CozE (CozEa) probed against EloR gave colorless spots of bacteria complying with no interaction between the two proteins. RodZ, MltG and YidC2 on the other hand gave blue spots when probed against EloR, suggesting that an interaction occurs. Positive and negative controls were supplied by the manufacturer (Euromedex). The Jag domain of EloR was tested against the cytosolic domain of MltG with and without the DUF domain. The interaction between the two domains were lost in the absence of the DUF domain in the cytosolic part of MltG. This indicates that the Jag domain of EloR interacts with the DUF domain of MltG.

Of all the proteins tested using BACTH, the positive hits were RodZ, YidC2 and MltG. MltG is a membrane protein predicted to be a lytic transglycosylase and is essential in *S. pneumoniae* (14). RodZ is, similar to EloR, considered to be part of the elongasome and studies in *E. coli* indicate that RodZ is important for the elongated cell shape (41). To test whether the interaction with these proteins were important for EloR localization, we wanted to analyze EloR-mKate2 localization in cells devoid of these genes. Localization of EloR-mKate2 in cells harboring deletions of either *rodZ* or *yidC2* (Figure 2B and 2C) shows that the midcell localization was observed in both mutants. Interestingly, however, in the genetic background lacking *yidC2*, we now detected an accumulation of EloR-mKate2 at the poles of the cells as well as at midcell. Further investigations of the polar localization of EloR-mKate2 revealed that the polar foci of EloR-mKate2 were found in old cellular poles (Supplemental Figure S4).

### EloR and MltG are part of the same complex

Since EloR was still localized at midcell when *rodZ* or *yidC2* were deleted, we hypothesized that MltG, which also has a midcell localization (15), could be important for this matter. However, MltG is essential in wild type cells making it impossible to track EloR-mKate2 in a Δ*mltG* mutant. Furthermore, we did not succeed in making an *mltG* depletion strain having EloR-mKate2 in the native *eloR* locus. Instead, to confirm the interaction between EloR and MltG *in vivo* in *S. pneumoniae* we attempted to use EloR as bait to pull down MltG. In order to do so, we constructed a mutant expressing a Flag-tagged EloR and an sfGFP-tagged MltG (strain AW447). By using resin beads tethered with α-Flag antibodies we pulled out Flag-EloR from the cell lysate as previously described by Stamsås et al., 2017. Then we looked for both Flag-EloR and sfGFP-MltG among the immunoprecipitated proteins using immunodetection and α-Flag and α-GFP antibodies (Figure 4). Indeed, when pulling out Flag-EloR using the α-Flag resin we found that sfGFP-MltG followed in the same fraction. Strain ds515 expressing only sfGFP-MltG was used as a negative control for a possible GFP/α-Flag interaction. In addition, to exclude a possible GFP/Flag-EloR unspecific interaction we co-expressed Flag-tagged EloR and GFP-tagged HlpA (DNA binding protein (42)) in strain AW459. When performing anti-Flag immunoprecipitation on lysates from this strain no HlpA-GFP was pulled down together with Flag-EloR.

**Figure 4.**
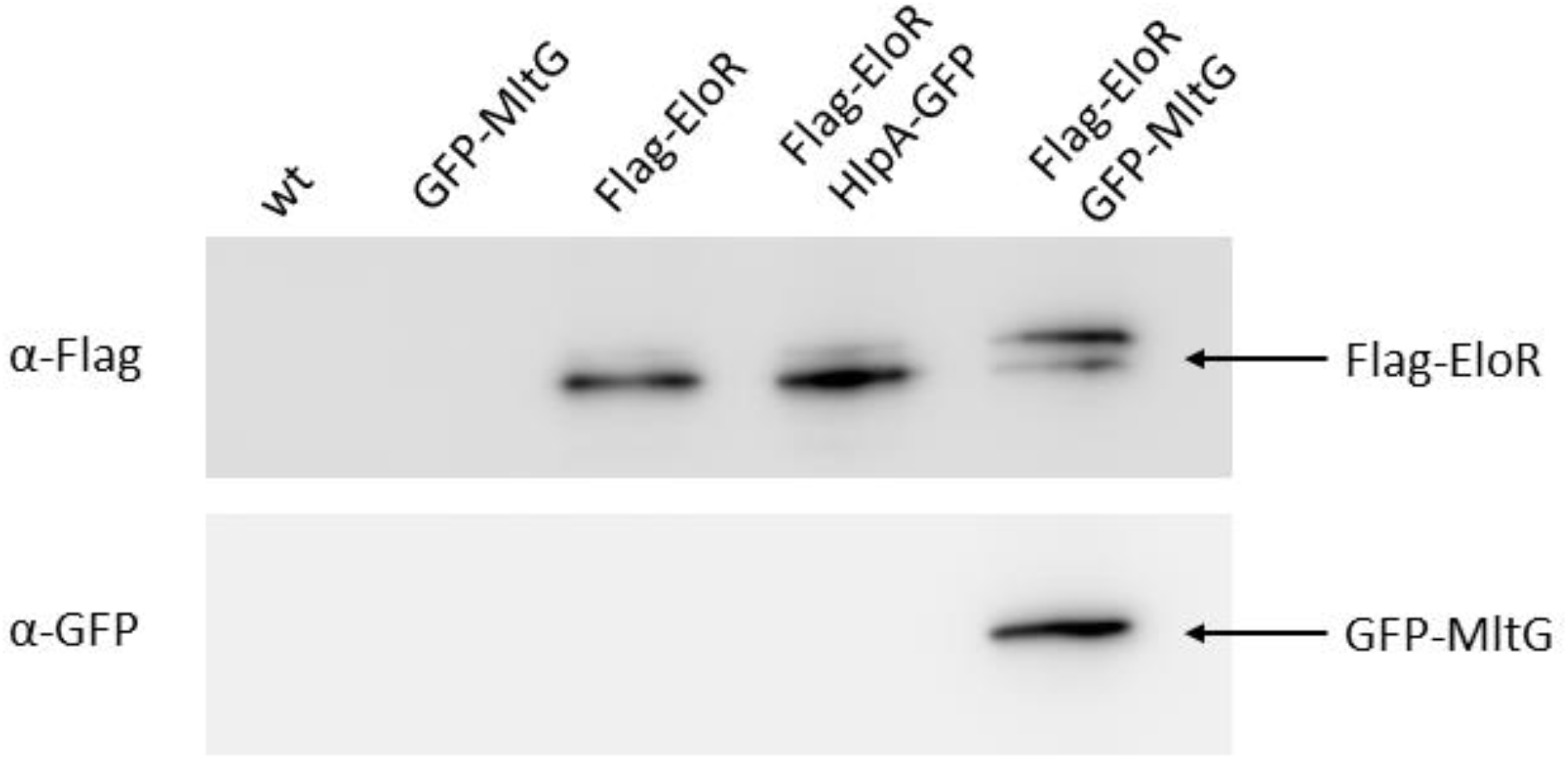
Immunoblot confirming the EloR – MltG interaction. Lysates from strains RH425 (wt), ds515 (*sfgfp-mltG*), AW98 (*flag-eloR*), AW459 (*flag-eloR, hlpA-gfp*), and AW447 (*flag-eloR, sfgfp-mltG*) were incubated with resin beads tethered with α-Flag antibodies to pull down Flag-EloR. As expected, immunoprecipitated Flag-EloR was found in strain AW98, AW459 and AW447, but not in strain ds515. sfGFP-MltG was only found in immunoprecipitated fractions when it was co-expressed with a Flag-tagged EloR.

### The Jag domain of EloR interacts with the intracellular DUF1346 domain of MltG

The immunoprecipitation result proved that EloR is in complex with MltG *in vivo* in *S. pneumoniae*. Our BACTH results suggested that the EloR/MltG interaction is direct. To further pinpoint the EloR-MltG interaction, we performed BACTH assays with the Jag domain of EloR (which is targeted to midcell) and the cytoplasmic part of MltG (Figure 5). Indeed, the sole Jag domain interacted with the cytosolic part of MltG (Figure 3). The cytosolic part of MltG mainly consists of a DUF-1346 (Domain of Unknown Function-1346) domain. When we test the cytosolic part of MltG lacking the DUF domain against the Jag domain of EloR, the interaction between the two are lost (Figure 3). Since MltG is located at the division zone of *S. pneumoniae* it is plausible that EloR is recruited to midcell through its interaction with MltG. Nevertheless, we cannot completely exclude the possibility that MltG is pulled down with EloR because both EloR and MltG interact with a common third protein. Further investigations (BACTH, co-IPs and cross-linking) are required to rule out this possibility.

**Figure 5.**
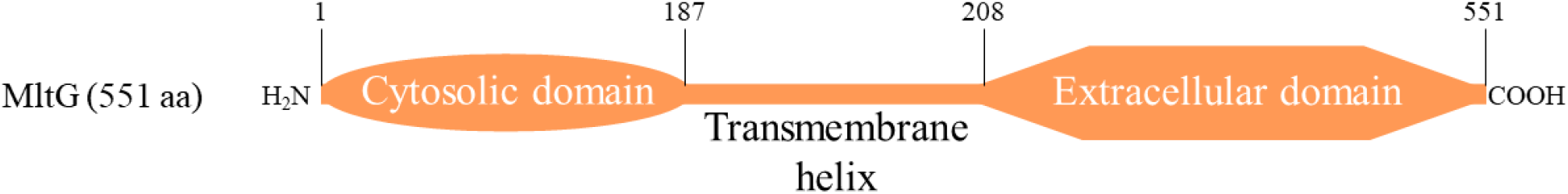
Schematic representation of MltG, including domain boarders.

### YidC2 does not affect MltG localization

Since EloR and MltG are part of the same complex, and since EloR localization was somewhat altered in the Δ*yidC2* mutant (localization to old poles, Figure 2C and Supplemental Figure S4), we wondered whether MltG, similar to EloR, would concentrate at the cell poles as well as at midcell in the Δ*yidC2* mutant. We therefore deleted *yidC2* in the strain expressing sfGFP-MltG from the native locus. However, like in wild-type cells, sfGFP-MltG was found at midcell in the Δ*yidC2* mutant, and no polar foci were observed (Supplemental figure S5A and S5B). Thus, in contrast to EloR, deletion of *yidC2* did not affect localization of MltG. This suggests that EloR may have additional interaction partners that need to be identified in the future, or possibly the RNA molecules it binds are concentrated at the old poles in the Δ*yidC2* mutant. The fact that EloR displayed interaction with YidC2 in BACTH assays and that absence of YidC2 induces altered EloR localization pattern suggest a functional role of YidC2 in the EloR/KhpA regulatory pathway. Since YidC proteins assist with insertion of membrane proteins during translation it is easy to imagine that the RNA-binding protein EloR is functionally linked to this process e.g. controlling the expression of one or several elongasome proteins. To unravel this would require additional research. The localization of sfGFP-MltG was not affected by the loss of *eloR*, as previously reported (19) and seen in Supplemental Figure S5C.

### Concluding remarks

It has previously been shown that knocking out the essential *pbp2b* results in suppressor mutations in *mltG, eloR* or *khpA* relieving the requirement of the elongasome in *S. pneumoniae* (15, 19). The current finding that EloR and MltG interact therefore corroborate that MltG and EloR are part of the same regulatory pathway. KhpA is also part of this complex since it has been shown previously that it interacts directly with EloR at the division zone (20, 21). In sum we can conclude that MltG, EloR and KhpA form a protein complex at the division zone, which regulates the elongasome on command from StkP. The relationship between EloR/KhpA and MltG is unknown. We have previously speculated that the expression levels of MltG is controlled though the RNA-binding capacity of EloR/KhpA. This, however, turned out to be wrong as the amount of MltG in a wild type background and in a Δ*eloR* mutant are similar (19). Another hypothesis is that the EloR/KhpA complex regulates the activity of MltG. MltG in *E. coli* have been shown to possess endolytic transglycosylase activity, i.e. breaking glycosidic bonds within a glycan strand (14, 43). Structural modelling and site directed mutagenesis of the active site of the pneumococcal MltG suggest that it has the same muralytic activity (15). It has been hypothesized that MltG in *S. pneumoniae* releases glycan strands made by both classes of PBPs so that they can be cross linked to new PG made by the divisome and elongasome (15, 43). In light of the recent discoveries regarding the function of class A PBPs, which suggest PBP1a to be involved in maturing the newly synthesized PG by filling in gaps, we propose a different model: MltG works together with amidases to open the PG layer so that PBP2b/RodA can add new PG to the existing layer and hence elongate the dividing cell (Figure 6). MltG must therefore be strictly regulated to avoid uncontrolled damage to the PG layer. The present study has shown just how important the MltG levels in the cells are: adjusting the expression level of MltG from an inducible promoter has proved to be difficult, and hence subsequent genetic changes to *eloR*, which is part of the same pathway, is not possible. Based on the data presented here the EloR/KhpA complex appears to play a role in regulation of MltG. Since inactivation of the RNA-binding domains of EloR gives the same phenotype as inactivation of the catalytic domain of MltG (PBP2b/RodA becomes redundant) (15, 19), one could speculate that the EloR/KhpA complex modulates the activity of MltG through RNA-binding. Another possibility is that EloR regulates the MltG activity directly through interaction with other cell division proteins. This must be confirmed or rejected by further experimental evidence.

**Figure 6.**
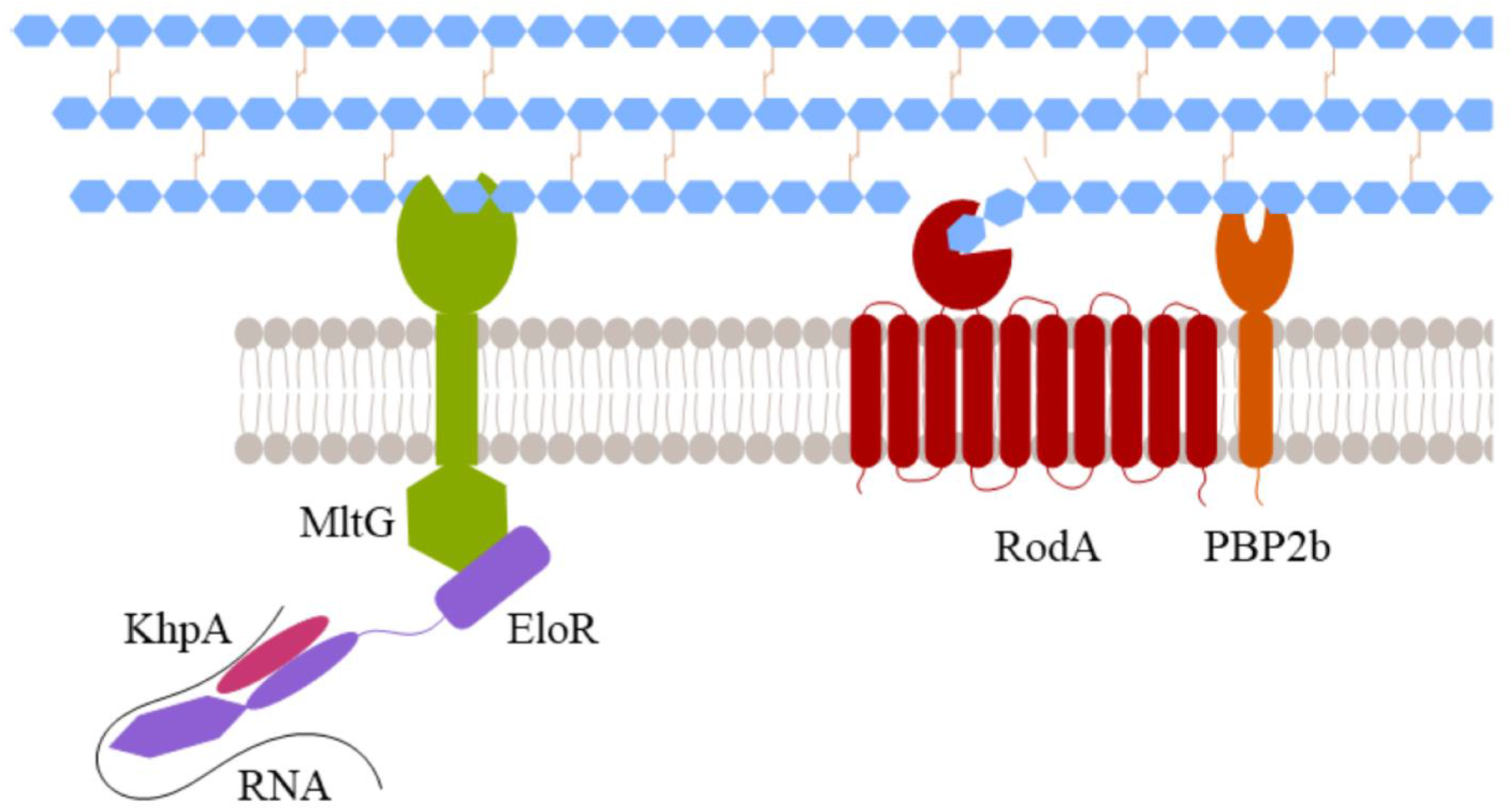
Modell of MltG/EloR/KhpA function. The communication between EloR/KhpA/RNA and MltG allows for a controlled opening of the PG. These openings are utilized by PBP2b/RodA to insert new PG in the lateral direction of the cell and in this way elongate the cell.

## Methods

### Bacterial strains, cultivation and transformation

All bacterial strains used in this work are listed in Table 1. All *E. coli* strains were grown in liquid LB broth with shaking or on LB agar plates at 30°C or 37°C. When necessary, the following antibiotic concentrations were used: 100 μg/ml ampicillin and 50 μg/ml kanamycin. Transformation of *E. coli* was performed with heat shock at 42°C for 30 seconds. All *S. pneumoniae* strains were grown in C-medium (44) without shaking or on Todd-Hewitt (TH) agar plates in an oxygen-depleted chamber using AnaeroGenTM bags from Oxoid at 37°C. Concentrations of 200 μg/ml streptomycin, 400 μg/ml kanamycin or 2 μg/ml chloramphenicol were employed when necessary. When introducing genetic changes, natural transformation was utilized. Exponentially growing cells were diluted to an OD_550_ of 0.05-0.1 and grown for two hours with 100-200 ng of the transforming DNA and 250 ng/ml CSP (final concentration) added to the growth medium. Thirty μl of the transformed cell cultures were plated on TH agar plates with the appropriate antibiotic and incubated at 37°C overnight.

**Table 1.** Bacterial strains and plasmids

| *S. pneumoniae*<br>strains | Relevant characteristics | Source |
| --- | --- | --- |
| R704 | R6 derivative, *comA::ermAM*; Ery<sup>r</sup> | J. P. Claverys |
| RH425 | R704, but streptomycin resistant; Ery<sup>r</sup>, Sm<sup>r</sup> | (48) |
| SPH131 | $\Delta comA$, P1::*P<sub>comR</sub>::comR*, *P<sub>comX</sub>::Janus*; Ery<sup>r</sup> Kan<sup>r</sup> | (35) |
| AW407 | $\Delta comA$, P1::*P<sub>comR</sub>::comR*, *P<sub>comX</sub>::eloR-mKate2*; Ery<sup>r</sup> Sm<sup>r</sup> | This work |
| AW408 | $\Delta comA$, P1::*P<sub>comR</sub>::comR*, *P<sub>comX</sub>::jag-mKate2*; Ery<sup>r</sup> Sm<sup>r</sup> | This work |
| AW409 | $\Delta comA$, P1::*P<sub>comR</sub>::comR*, *P<sub>comX</sub>::jag-linker-mKate2*; Ery<sup>r</sup><br>Sm<sup>r</sup> | This work |
| AW410 | $\Delta comA$, P1::*P<sub>comR</sub>::comR*, *P<sub>comX</sub>::linker-mKate2*; Ery<sup>r</sup> Sm<sup>r</sup> | This work |
| AW420 | $\Delta comA$, P1::*P<sub>comR</sub>::comR*, *P<sub>comX</sub>::eloR<sup>K36A</sup>-mKate2*; Ery<sup>r</sup>,<br>Sm<sup>r</sup> | This work |
| AW424 | $\Delta comA$, P1::*P<sub>comR</sub>::comR*, *P<sub>comX</sub>::eloR<sup>K37A</sup>-mKate2*; Ery<sup>r</sup>,<br>Sm<sup>r</sup> | This work |
| AW425 | $\Delta comA$, P1::*P<sub>comR</sub>::comR*, *P<sub>comX</sub>::eloR<sup>F39A</sup>-mKate2*; Ery<sup>r</sup>,<br>Sm<sup>r</sup> | This work |
| AW426 | $\Delta comA$, P1::*P<sub>comR</sub>::comR*, *P<sub>comX</sub>::eloR<sup>L40M</sup>-mKate2*; Ery<sup>r</sup>,<br>Sm<sup>r</sup> | This work |
| AW453 | $\Delta comA$, P1::P<sub>comR</sub>::comR, P<sub>comX</sub>::eloR-mKate2,<br>$\Delta stkP$::janus; Ery<sup>r</sup>, Km<sup>r</sup> | This work |
| AW415 | $\Delta comA$, P1::P<sub>comR</sub>::comR, P<sub>comX</sub>::eloR-mKate2,<br>$\Delta yidC2$::janus; Ery<sup>r</sup>, Km<sup>r</sup> | This work |
| AW417 | $\Delta comA$, P1::P<sub>comR</sub>::comR, P<sub>comX</sub>::eloR-mKate2,<br>$\Delta rodZ$::janus; Ery<sup>r</sup>, Km<sup>r</sup> | This work |
| AW447 | $\Delta comA$, m(sf)gfp-mltG, flag-eloR; Ery<sup>r</sup> Sm<sup>r</sup> | This work |
| AW459 | $\Delta comA$, flag-eloR, hlpA-gfp-chloramphenicol; Ery<sup>r</sup>, Sm<sup>r</sup>,<br>Cam<sup>r</sup> | This work |
| DS515 | $\Delta comA$, m(sf)gfp-mltG; Ery<sup>r</sup>, Sm<sup>r</sup> | (19) |
| AW98 | $\Delta comA$, flag-eloR; Ery<sup>r</sup>, Sm<sup>r</sup> | (19) |
| MH43 | $\Delta comA$, m(sf)gfp-mltG; $\Delta yidC2$::janus; Ery<sup>r</sup>, Km<sup>r</sup> | This work |
| SPH472 | $\Delta comA$, m(sf)gfp-mltG; $\Delta eloR$::janus; Ery<sup>r</sup>, Km<sup>r</sup> | (19) |
| ***E. coli* strains** |  |  |
| XL1-Blue | Host strain | Agilent<br>Technologies |
| BTH101 | BACTH expression strain, cya- | Euromedex |
| **Plasmids** |  |  |
| pKT25 | Plasmid used in BACTH analysis | Euromedex |
| pKNT25 | Plasmid used in BACTH analysis | Euromedex |
| pUT18C | Plasmid used in BACTH analysis | Euromedex |
| pUT18 | Plasmid used in BACTH analysis | Euromedex |
| pKNT25-eloR | T25 domain fused to the C-terminus of EloR | (19) |
| pKT25-jag | T25 domain fused to the N-terminus of the Jag domain of EloR | This work |
| pUT18C-mltG | T18 domain fused to the N-terminus of MltG | (19) |
| pUT18C-mltG<sub>cyt</sub> | T18 domain fused to the N-terminus of the cytoplasmic domain of MltG | This work |
| pUT18C-mltG<sub>cyt</sub>ΔDUF | T18 domain fused to the N-terminus of the cytoplasmic domain of MltG without DUF | This work |
| pUT18C-pbp2b | T18 domain fused to the N-terminus of PBP2b | (49) |
| pUT18C-rodA | T18 domain fused to the N-terminus of RodA | (49) |
| pUT18C-rodZ | T18 domain fused to the N-terminus of RodZ | (19) |
| pUT18C-mreC | T18 domain fused to the N-terminus of MreC | (19) |
| pUT18C-mreD | T18 domain fused to the N-terminus of MreD | This work |
| pUT18-cozE | T18 domain fused to the C-terminus of CozE | (49) |
| pUT18-yidC2 | T18 domain fused to the C-terminus of YidC2 | This work |

### DNA constructs

All primers used in this study are listed in Table 2. DNA constructs used to transform *S. pneumoniae* were made using overlap extension PCR (45). In short, in order to create deletion mutants, approximately 1000 bp sequence upstream and downstream of the gene in question were amplified and fused with the 5’ end and 3’ end of the Janus cassette (46), respectively. The same flanking regions were then used to replace the Janus cassette with an alternative DNA sequence (47). Constructs used to produce BACTH plasmids were amplified from *S. pneumoniae*, cleaved with restriction enzymes (XbaI and EcoRI from New England BioLabs), and ligated into the preferred plasmid using Quick ligase (New England BioLabs). The plasmids used in this study are listed in Table 1. All constructs were verified with DNA sequencing.

**Table 2.**
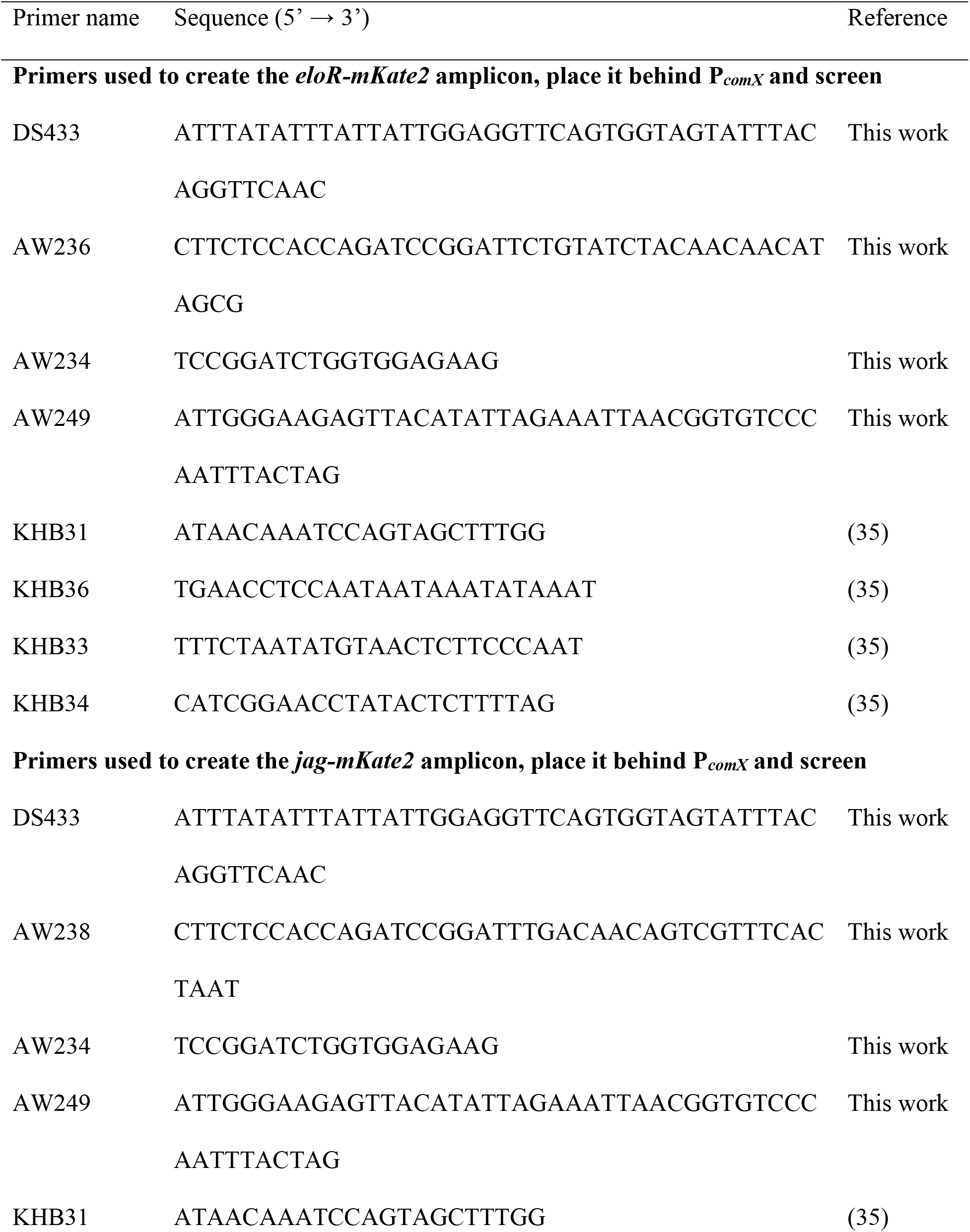
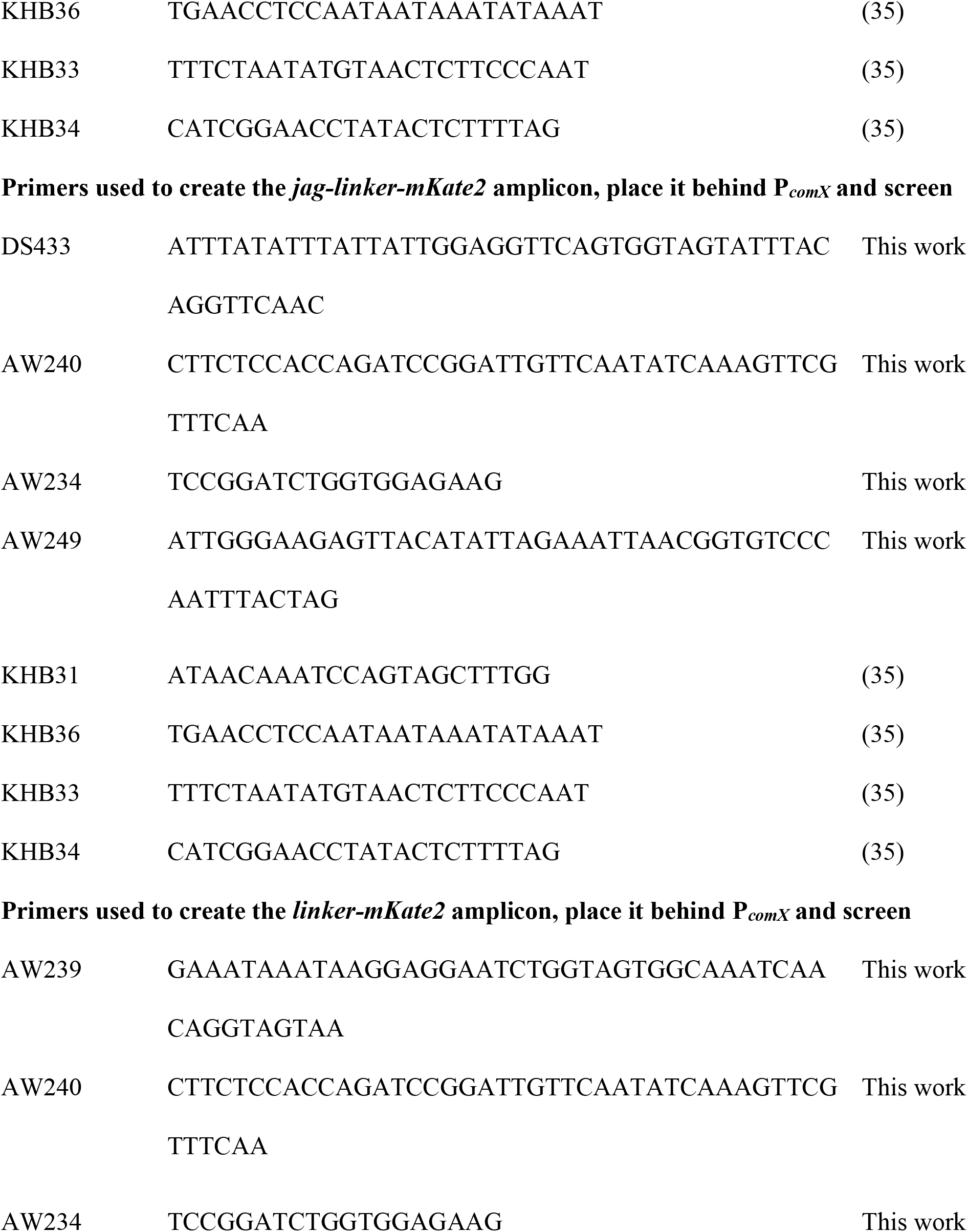
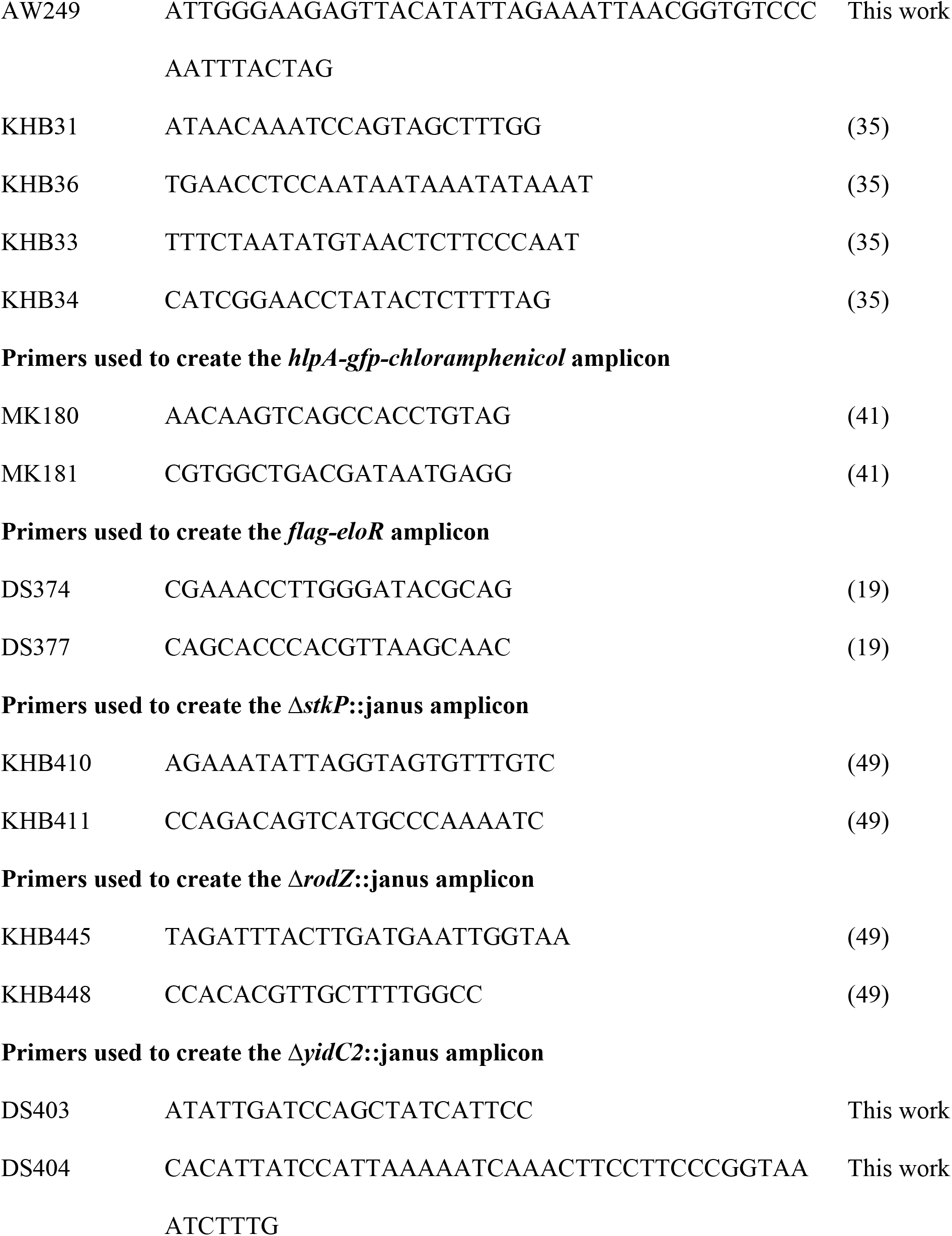
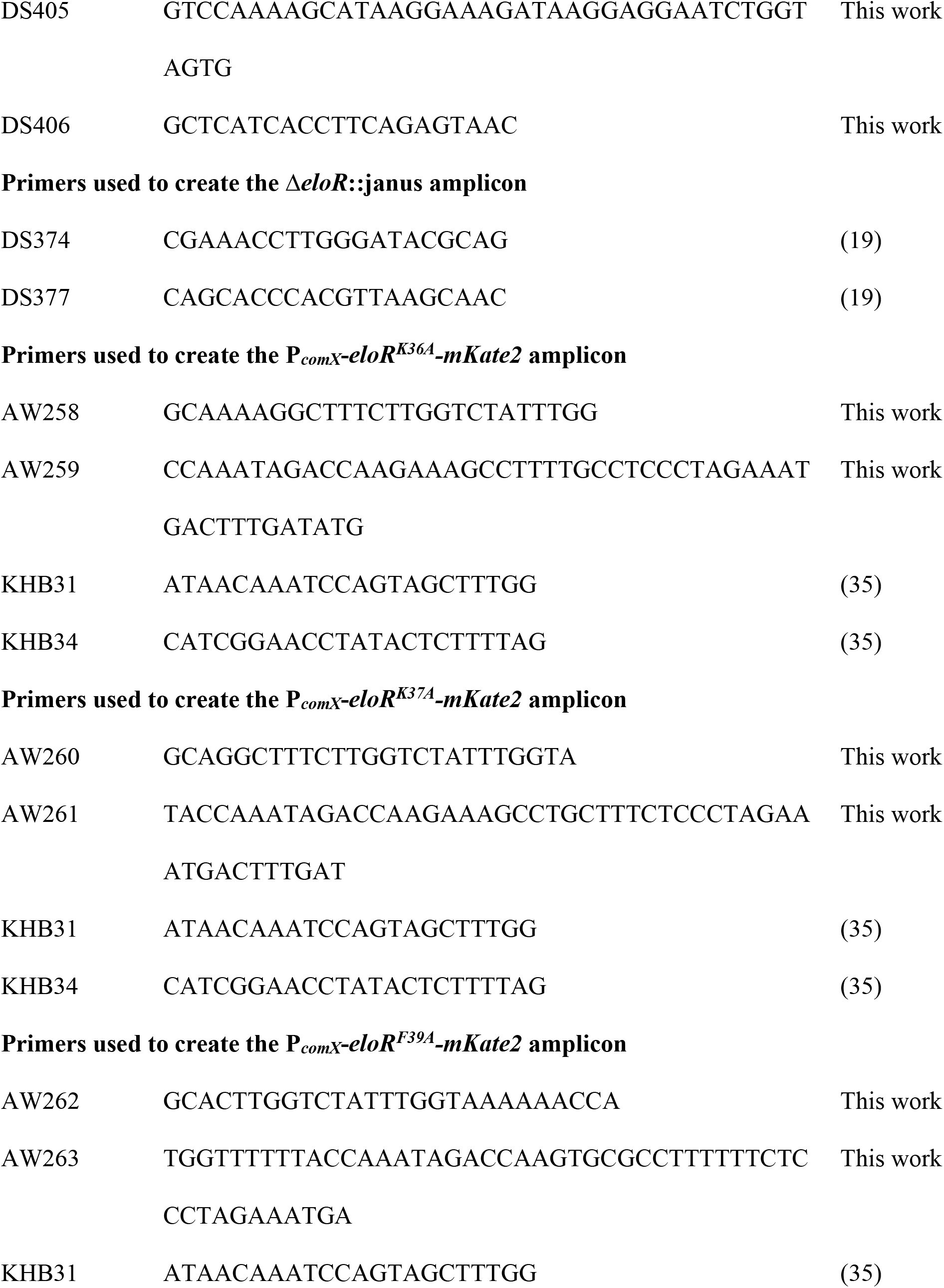
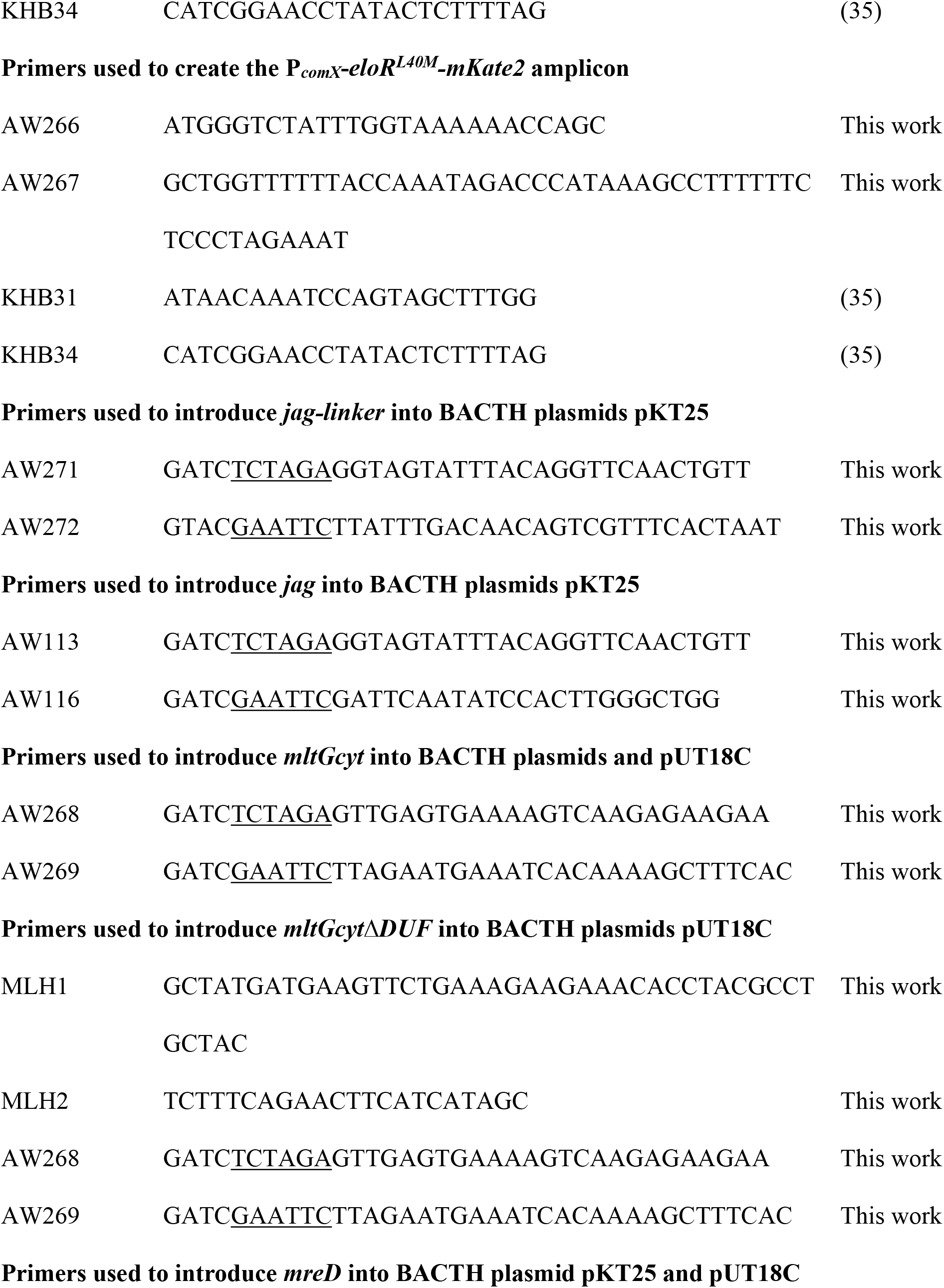
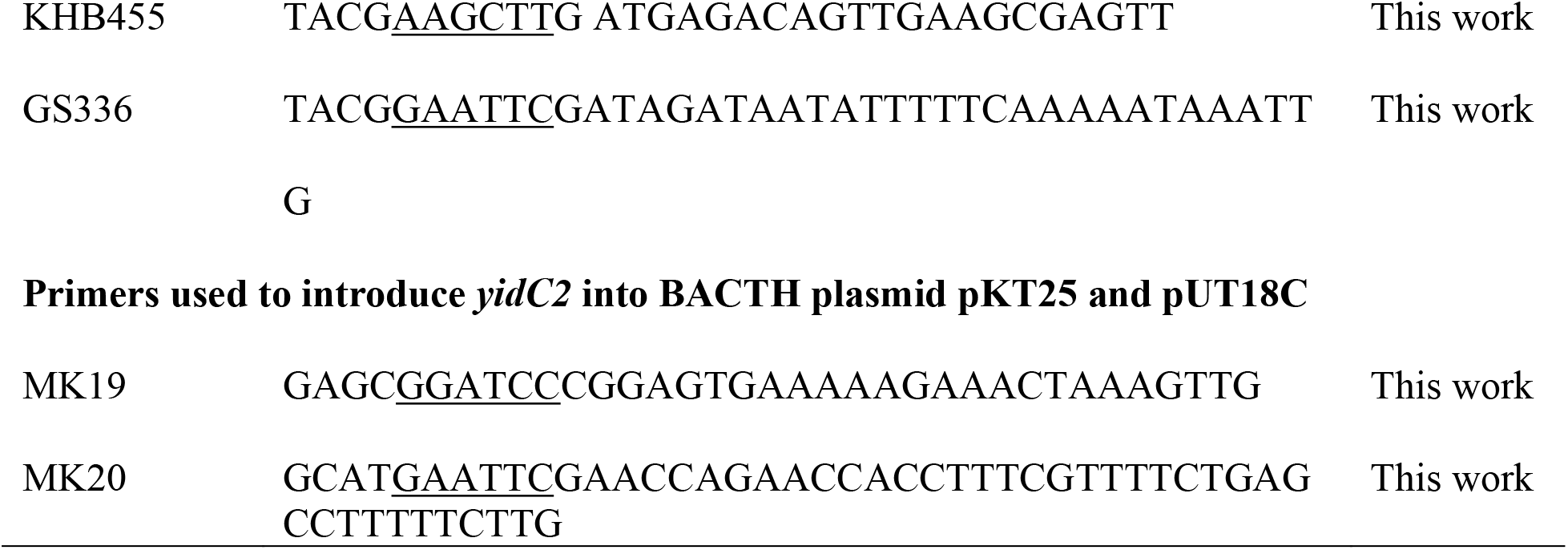
Primers

### Bacterial two hybrid assay

Bacterial two hybrid (BACTH) assays are based on the two tags T18 and T25 that make up the catalytic domain of *Bordetella pertussis* adenylate cyclase (CyaA). In order to test whether two proteins interact, their genes are cloned in frame with one tag each and co-expressed in *E. coli* BTH101 cells (*cyaA-*). If the two proteins interact, T18 and T25 are brought into close proximity to make up an active CyaA catalytic domain. This results in cAMP production which induces expression of *lacZ* (β-galactosidase). β-galactosidase cleaves X-gal, resulting in blue bacteria on X-gal containing agar plates. In instances where the two tested proteins do not interact, no β-galactosidase is expressed, and the bacteria remain white. The BACTH experiments were performed as described by the manufacturer (Euromedex). The genes encoding our proteins of interest were cloned in reading frame with either the T18 or T25 encoding gene in the plasmids pUT18, pUT18C, pKNT25 and pKT25. The plasmids were then transformed into *E. coli* XL1-Blue cells, then isolated and sequenced. In order to test the interaction between two proteins, they were co-expressed with one tag (T18, T25) each in *E. coli* BTH101 cells. After overnight incubation of transformants, five random colonies were picked, grown to exponential phase, and spotted (2 μl) onto LB agar plates containing ampicillin (100 μg/ml), kanamycin (50 μg/ml), IPTG (0.5 mM) and X-gal (40μg/ml). After overnight incubation at 30°C the results were documented.

### Co-immunoprecipitation and western blotting

Co-IP was performed using ANTI-FLAG^®^ M2 affinity gel (Sigma-Aldrich). *S. pneumoniae* strains were grown to OD_550_ = 0.3 and lysed with 1 ml lysis buffer (50 mM Tris HCl, pH 7.4, 150 mM NaCl, 1 mM EDTA, 1% Triton X-100) by triggering the endogenous pneumococcal autolysin LytA at 37°C for 5 minutes. The lysate was incubated with 40 μl ANTI-FLAG^®^ M2 affinity gel with gentle rotation at 4°C overnight. After washing the affinity gel three times with 500 μl TBS, SDS sample buffer was added and the samples were incubated at 95°C for 10 minutes. Proteins from eight μl of each sample were separated in a 12 % SDS PAGE gel. After electrophoresis the separated proteins were electroblotted onto a PVDF membrane using a Trans-Blot Turbo Transfer System (Bio-Rad) with a standard protocol for seven minutes. Finally, Flag-tagged proteins were detected as previously described by Stamsås et al, 2017. GFP-tagged proteins were detected with Chromotek rabbit polyclonal antibody for GFP, using the same protocol as above and dilutions as recommended by the manufacturer.

### Phase contrast and fluorescent microscopy

Cells were prepared for microscopic imaging by growing them to OD_550_ = 0.4, then diluting the culture to OD_550_ = 0.1 and grown for another hour prior to microscopy. When relevant, 2 μM ComS inducer was added. Proteins fused with the fluorescent mKate2 were visualized as previously described (19) using a Zeiss AxioObserver with ZEN Blue software, an ORCA-Flash 4.0 V2 Digital CMOS camera (Hamamatsu Photonics), and a 100x phase-contrast objective. An HXP 120 Illuminator (Zeiss) was used as a fluorescence light source. Images were prepared and analyzed using the ImageJ software with the MicrobeJ plugin.

