## Supplemental info for "EloR interacts with the lytic transglycosylase MltG at midcell in *Streptococcus pneumoniae* R6"

**Supplementary information**

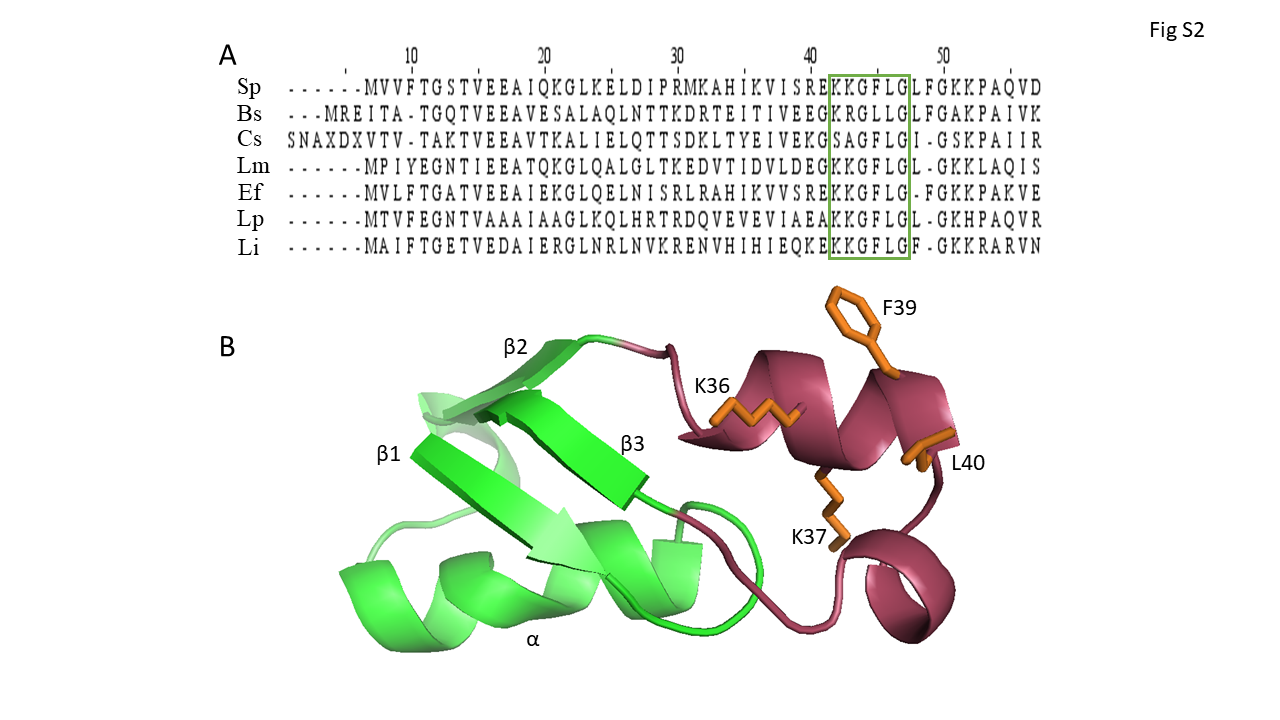

**Figure S1.** A) Alignment of the Jag domains from (listed in same order as they appear in image) *S. pneumoniae*, *B. subtilis*, *C. symbiosium*, *Listeria monocytogenes*, *Enterococcus faecalis*, *Lactobacillus plantarum*, and *Lactococcus lactis*. The conserved KKGFLG (green box) is indicated. B) Predicted structure (iTasser) of the Jag domain of *S. pneumoniae* EloR. The β-α-β-β fold with the α-helix laying on top of a three-stranded β-sheet is portrayed in green. The loop connecting the second and third β-strand is portrayed in red. The KKGFLG motif is shown in orange sticks.

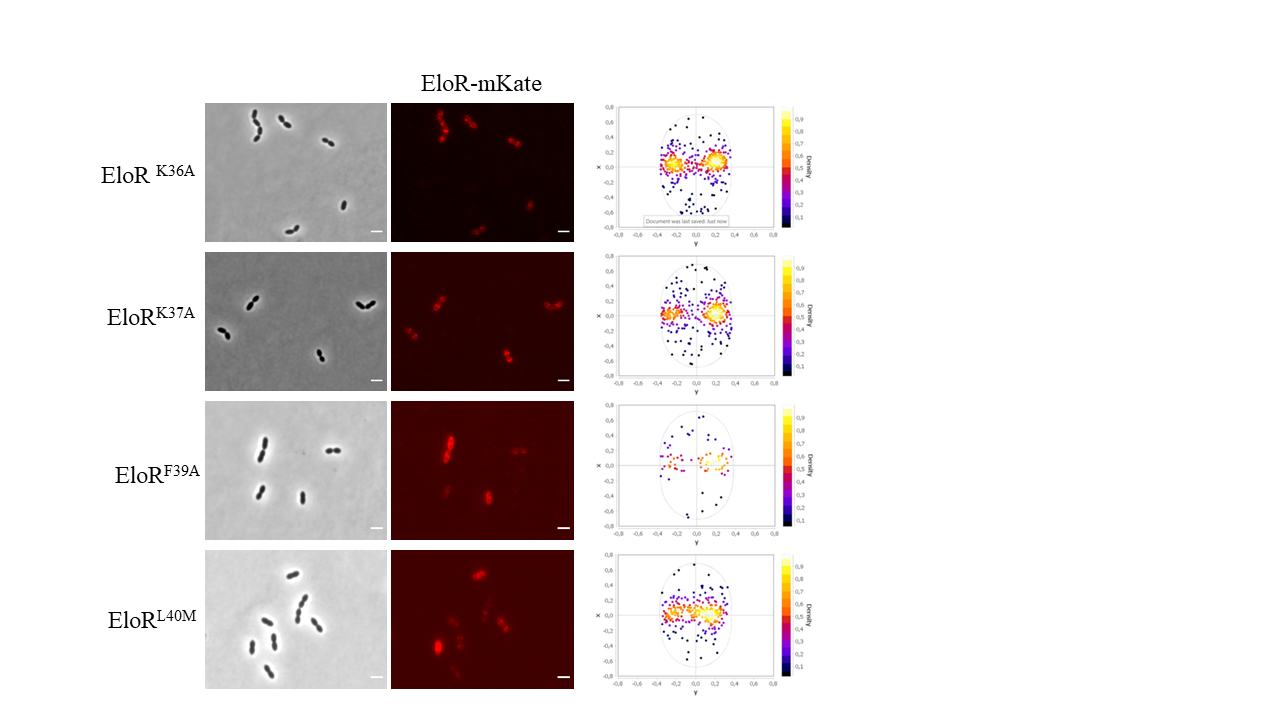

**Figure S2.** Localization of EloR-mKate2 harboring the amino acid substitutions K36A, K37A, F39A and L40M. EloR-mKate2 is found concentrated at midcell with all the introduced mutations, as confirmed by the fluorescent maximum signals. The number of cells analyzed were N = 188 for K36A, N = 183 for K37A, N = 182 for F39A and N = 189 for L40M. Scale bars are 2 µm.

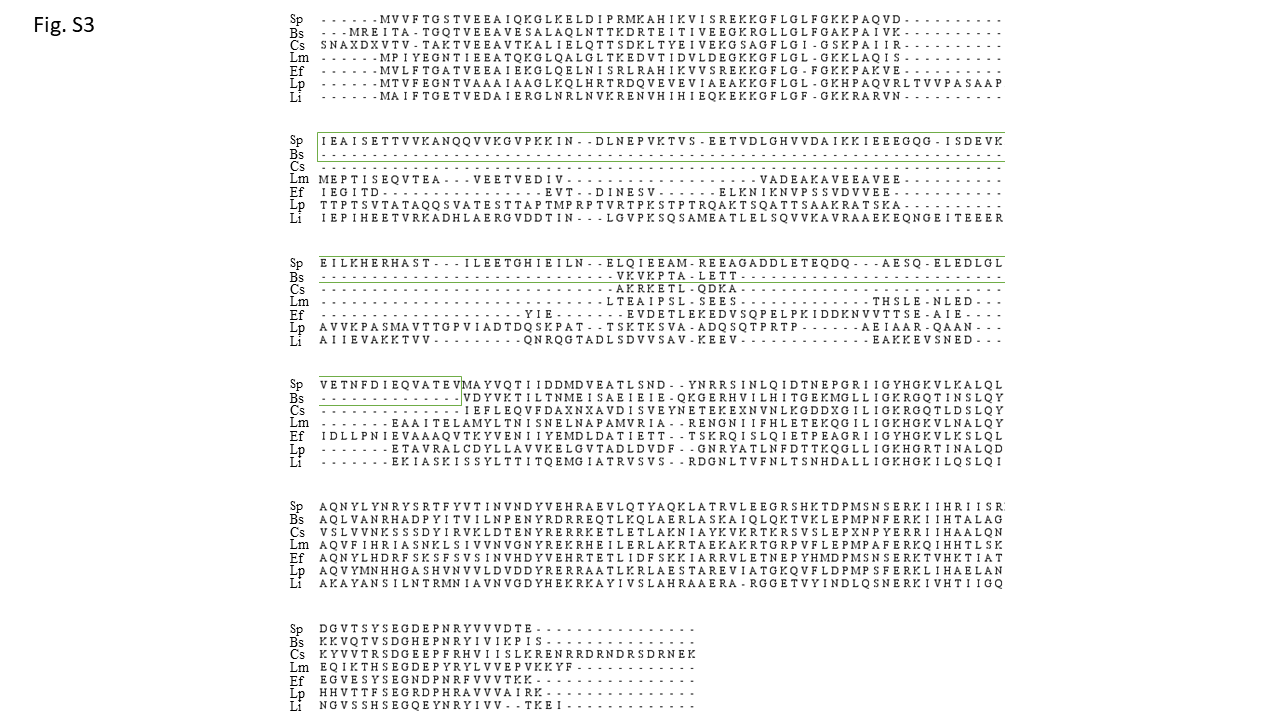

**Figure S3.** Alignment of EloR from (listed in same order as they appear in image) *S. pneumoniae*, *B. subtilis*, *C. symbiosium*, *Listeria monocytogenes*, *Enterococcus faecalis*, *Lactobacillus plantarum*, and *Lactococcus lactis*. The linker domain of EloR from *S. pneumoniae* (green box) is made up of approximately 135 amino acids, while the equivalent domain from *B. subtilis* (green box) only consists of approximately 10 amino acids.

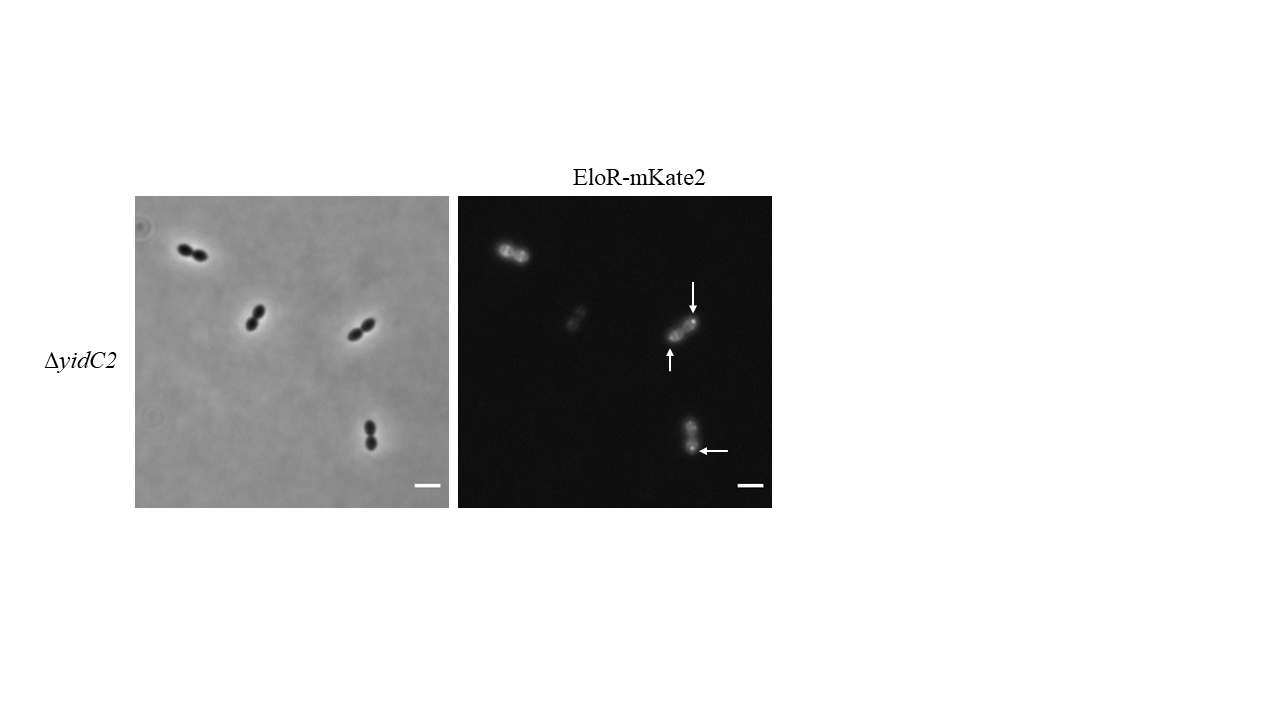

**Figure S4.** Polar localized EloR-mKate2 in a ∆*yidC2* mutant is typically found in old cell poles as seen indicated by white arrows. Scale bars are 2 µm.

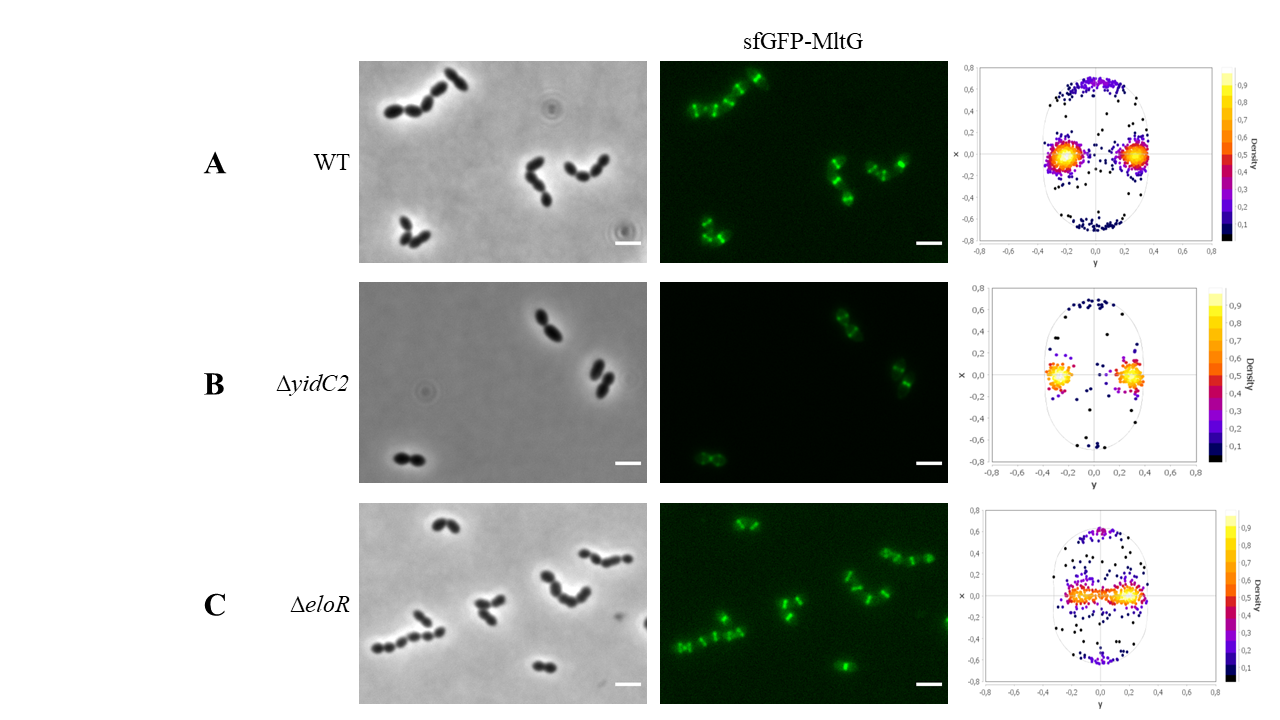

**Figure S5.** Localization of sfGFP-MltG in a A) wild type background (N=1209), B) ∆*yidC2* mutant (N=253), and C) ∆*eloR* mutant (N=465). sfGFP-MltG was found localized at midcell in all genetic backgrounds investigated. Scale bars are 2 µm.
